# Pathways of inter- and intrasubunit allosteric signaling in the C-linker disk and cyclic nucleotide-binding domain of HCN2 channels

**DOI:** 10.1101/2020.06.13.150086

**Authors:** Christopher Pfleger, Jana Kusch, Mahesh Kondapuram, Tina Schwabe, Christian Sattler, Klaus Benndorf, Holger Gohlke

## Abstract

Opening of hyperpolarization-activated cyclic nucleotide-modulated (HCN) channels is controlled by membrane hyperpolarization and binding of cyclic nucleotides to the tetrameric cyclic nucleotide-binding domain (CNBD), attached to the C-linker disk (CL). Confocal patch-clamp fluorometry revealed a pronounced cooperativity of ligand binding among protomers. However, by which pathways allosteric signal transmission occurs remained elusive. Here, we investigate how changes in the structural dynamics of the CL-CNBD of mouse HCN2 upon cAMP binding relate to inter- and intrasubunit signal transmission. Applying a rigidity theory-based approach, we identify two intersubunit and one intrasubunit pathways that differ in allosteric coupling strength between cAMP binding sites or towards the CL. These predictions agree with results from electrophysiological and patch-clamp fluorometry experiments. Our results map out distinct routes within the CL-CNBD that modulate different cAMP binding responses in HCN2 channels. They signify that functionally relevant submodules may exist within and across structurally discernable subunits in HCN channels.

## Introduction

Hyperpolarization-activated cyclic nucleotide-modulated (HCN) channels belong to the superfamily of voltage-gated ion channels and mediate electrical pacemaking activity in specialized cells of the heart and brain (1). The crucial role of hyperpolarization-activated currents makes these channels promising drug targets (1, 2). Four human isoforms (HCN1 – HCN4) are known, which all form functional homo- and heterotetrameric channels (3). Recently, the first full-length 3D structures of the human HCN1 (4) and HCN4 (PDB id 6gyo and 6gyn) channels in the presence and absence of cAMP were resolved. Each subunit consists of four domains, the transmembrane domain (TMD) composed of six transmembrane helices, comprising the voltage sensor and the channel pore (4), the N-terminal HCN domain, and the C-terminal cyclic nucleotide-binding domain (CNBD), which is connected to the TMD via the C-linker (CL) domain (Figure 1A, B).

**Figure 1:**
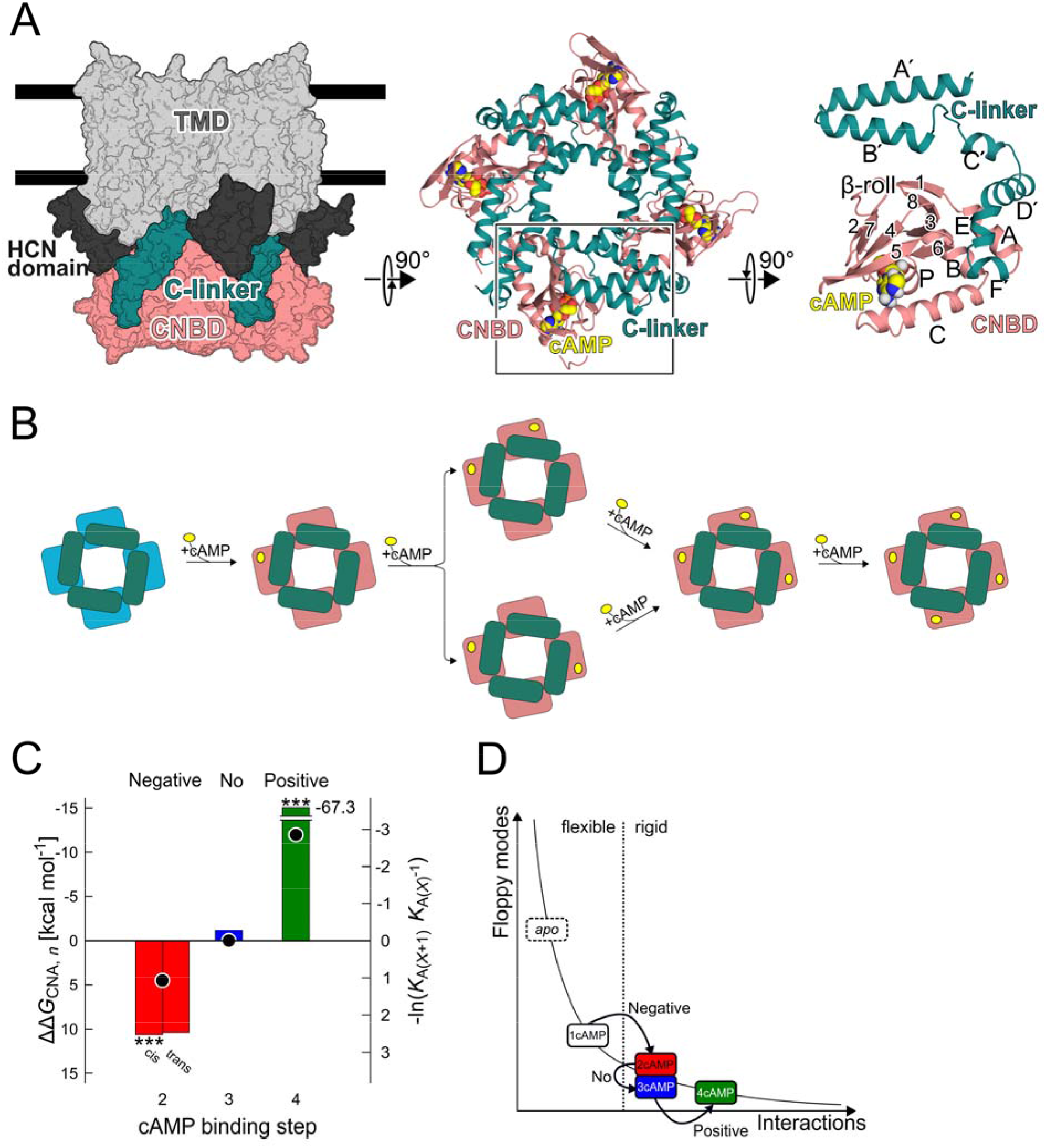
Changes in structural rigidity of the CL-CNBD underlie the complex succession of cooperativities for cAMP binding steps. (*A*) Schematic representation of the tetrameric HCN2 channel depicting its domain organization: transmembrane domain (TMD, grey), HCN domains (black), C-linker (CL, cyan) region, and cyclic nucleotide-binding domain (CNBD, salmon). The CL connects the CNBD to the TMD. Each CL-CNBD is composed of helices A-C, P and strands 1-8 (β-roll) in the CNBD and helices A’-F’ in the CL region; bound cAMP is indicated in yellow. The structure is taken from PDB ID 1q5o. (*B*) Illustration of the sequence of cAMP binding to the tetrameric CNBD with the two possible cAMP configurations (*cis*/*trans*) at the second binding step. (*C*) Left y-axis and bars: ΔΔ*G*_*CNA, n*_ (eq. 1) quantifying the non-independency among cAMP binding processes; stars denote values significantly different from zero (*n* = 10, *: *p* < 0.05, **: *p* < 0.01, ***: *p* < 0.001); the terms “negative”, “no”, and “positive” relate to associated cooperativities in the binding events. Right y-axis and dots: Microscopic association constants of cAMP binding to non-activated HCN2 channels and derived succession of cooperativities “negative – no – positive”; values were taken from ref. (13). (*D*) Illustration of the occurrence of negative, no, and positive cooperativity of cAMP binding from entropy changes due to altered structural rigidity (“floppy modes” denote internal independent degrees of freedom). The vertical dashed line indicates the rigidity percolation threshold where biomolecules switch from a flexible to a rigid state. See ref. (20) for further details.

HCN channels are dually regulated (5, 6) by membrane hyperpolarization and the binding of cyclic nucleotides (cNMP) to the CNBD. cAMP is the physiological agonist for HCN2 channels (5). According to the tetrameric quaternary structure of HCN channels, up to four cNMPs can bind to a channel (Figure 1B). Binding of cNMP to the CNBD stabilizes a state that fosters the opening of HCN channels by disinhibition (7, 8). That way, both the rate and the extent of channel activation evoked by hyperpolarization are increased, and steady-state activation is shifted to less negative voltages (5, 9). The underlying molecular mechanisms of how both stimuli activate HCN channels are still poorly understood.

Results from confocal patch-clamp fluorometry, a method allowing to record cNMP binding and activation gating in parallel (10, 11), revealed a 3-fold increase of the overall cAMP affinity upon HCN2 channel activation, proving thus the principle of reciprocity between ligand binding and activation in HCN2 channels (12). This approach yielded furthermore microscopic association constants that suggest a pronounced cooperativity among the four subunits, with a succession “positive-negative-positive” for the second, third, and fourth cAMP binding step for activated HCN2 channels and “negative-no-positive” for non-activated ones (13). However, by which pathways the allosteric signal transmission occurs between subunits of the CL-CNBD and/or towards the TM core remained elusive.

Structural comparisons of the full-length HCN1 and HCN4 channels in the presence and absence of cAMP reveal only subtle differences within the CL-CNBD domains, with the largest conformational differences found for the terminal helix C within the CNBD. In the presence of cAMP, the tetrameric CL-CNBD arrangements also globally rotate relative to the TM domains (4). Upon cAMP binding, earlier tmFRET and DEER studies on isolated CL-CNBDs revealed, besides changes in the location of maxima of distance distributions, indicative of small conformational changes, changes in the width of such distributions (14-17), indicative of a decrease of the number of accessible configurational states of the CL-CNBD. Together, these results point to changes in the channel dynamics underlying the allosteric signaling within the CL-CNBD. In the extreme case, allosteric signaling can appear even in the absence of conformational changes (18): the allosteric communication then arises from changes in frequencies and amplitudes of biomolecular thermal fluctuations, and this “dynamic allostery” is primarily an entropic effect (19).

Here, we address at the atomistic level the question how changes in the structural dynamics of the CL-CNBD of mouse HCN2 (mHCN2) channels upon cAMP binding relate to inter- and intrasubunit allosteric signal transmission. Applying an ensemble- and rigidity theory-based free energy perturbation approach to analyze dynamic allostery (20), we identify two intersubunit pathways and one intrasubunit pathway of allosteric signal transmission that differ in the strength of allosteric coupling between cAMP binding sites or towards the CL region. These predictions are concordant with results from electrophysiological and patch-clamp fluorometry experiments on wildtype and mutant mHCN2. Our results map out distinct routes within the CL-CNBD that modulate different cAMP binding responses in HCN2 channels through disjunct salt bridge interactions.

## Results

### Non-uniform changes in structural rigidity of the CL-CNBD underlie the complex succession of cooperativities for cAMP binding steps

As a model system for studying the subunit cooperativity, we used the fully cAMP-bound X-ray crystal structure of the isolated tetrameric CL-CNBD from mHCN2 in its inactive state. From this structure, all possible cAMP-bound structures, with one, two (*cis* and *trans* configuration), and three cAMP molecules, were generated by removing respective cAMP(s). The resulting structures served as input for ten independent MD simulations of 1 μs length each, yielding ensembles with 2,000 conformations each. We then analyzed the ensembles using our rigidity theory-based free energy perturbation approach (20). In this approach, biomolecules, including ligands, are modeled as constraint networks where nodes represent the atoms and constraints the covalent and non-covalent binding interactions. Such a constraint network is efficiently decomposed into rigid clusters and connecting flexible hinge regions by applying rigidity theory (21). Biomolecules generally display a hierarchy of structural stability that reflects the modularity of their structure (22). To identify this hierarchy, a “constraint dilution trajectory” of network states, obtained from an initial network topology by successively removing non-covalent constraints, is analyzed (see Materials and Methods) (22-26). The networks of the generated conformations were perturbed by removing all covalent and non-covalent constraints associated with cAMP(s) to compare changes in biomolecular structural stability between the ground (cAMP-bound) and perturbed (unbound) state. Note that upon perturbation, the conformation of the protein remains unchanged. The results were then post-processed to predict the type of cooperativity associated with each cAMP binding step (20). To do so, we computed the sum of per-residue free energies Δ*G*_CNA_ associated with the change in biomolecular structural stability due to the removal of (a) ligand(s) for all cAMP-bound states (Δ*G*_*CNA, 1cAMP*_, Δ*G*_*CNA, 2cAMP/cis*_, Δ*G*_*CNA, 2cAMP/trans*_, Δ*G*_*CNA, 3cAMP*_, Δ*G*_*CNA, 4cAMP*_) (eq. 4, Table S1). From that, ΔΔ*G*_*CNA,n*_ was computed (eq. 1; Figure 1C)

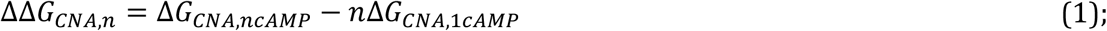

which quantifies the non-independency among binding processes; Δ*G*_*CNA,ncAMP*_ is the free energy associated with cAMP binding to *n* binding sites, and *n* Δ*G*_*CNA,1cAMP*_ provides a normalization in terms of *n*-fold binding of one cAMP. This definition allows for comparing our results to ref. (13), where a global-fit strategy involving a Markovian state model with four binding steps provided a full set of microscopic association constants *K*_AX_ for all cAMP binding steps to the non-activated HCN2 channel; cooperativity of cAMP binding there was quantified from successive *K*_AX_ (Figure 1C).

The computed ΔΔ*G*_*CNA,n*_ values display a succession of positive, zero, and negative values for the binding of two, three, and four cAMP to tetrameric CL-CNBD in the non-activated conformation, respectively (Figure 1C), associated with negative, no, and positive cooperativity in the binding events. This result is in remarkable agreement with the experiment (Figure 1C) (13), which thereby justifies the applicability of the rigidity theory-based free energy perturbation approach for this system. The existence of a heterogeneous succession of cooperativity among identical subunits has also been reported for other circular biomolecules (27). For the binding of two cAMP molecules, ΔΔ*G*_*CNA*,2_ is 10.6 ± 2.06 and 10.4 ± 4.84 kcal mol^−1^ for *cis* and *trans* configurations, respectively. Both values are not significantly different, which is in agreement with previous results that demonstrated similar ligand-evoked activating effects of subunits for the *cis* and the *trans* position in HCN2 constructs with only two functional binding sites (28), but at variance with other results showing functional differences between the *cis* and *trans* configuration (29). The magnitude of ΔΔ*G*_*CNA,*4_ in the case of binding of four cAMP is the largest, qualitatively concordant with experiment (Figure 1C) (13) and indicating a very strong positive cooperativity involved in reaching a fully bound CL-CNBD.

These results demonstrate a sequential, but non-uniform loss of independent internal degrees of freedom of the CL-CNBD upon cAMP binding steps (Figure 1D, Figure S1) in the context of rigidity percolation. At the rigidity percolation threshold, a phase transition occurs at which the network loses (almost) all of its independent internal degrees of freedom, and rigidity starts to percolate through the network (26, 30). At the threshold, the network is most sensitive to changes, i.e., adding or removing a few constraints, e.g., due to ligand binding, causes a rigid or flexible state. Away from the threshold, additional or fewer constraints have no or only minor effects on network stability (31).

The existence of intersubunit cooperativity has been suggested to be a general feature of circular biomolecules built by identical allosteric protomers (32). Such biomolecules can act as a switch, in which the system changes from a fully inactive to a fully active state depending on the number of ligand-bound subunits (32). In conjunction with our model of cooperativity, the relatively low ΔΔ*G*_*CNA,n*_ values for ensembles with two or three cAMP-bound CL-CNBD subunits show a pre-stabilized tetrameric CL-CNBD structure. Further interactions due to the presence of the fourth cAMPs then yield a super-additive effect in structural stabilization of the CL-CNBDs, resulting in the large ΔΔ*G*_*CNA,*4_. This extra stabilization due to the fully liganded state particularly involves the helices B and C within the CNBD, even in subunits to which cAMP was bound already (Figure S2).

To conclude, the computed ΔΔ*G*_*CNA,n*_ values display a succession of positive, zero, and negative values for the binding of two, three, and four cAMP molecules to the tetrameric CL-CNBD in the non-activated conformation, respectively, associated with negative, no, and positive cooperativity in the binding events, in agreement with the experiments. These results demonstrate that non-uniform changes in structural rigidity of the CL-CNBD underlie the complex succession of cooperativities for cAMP binding steps.

### Changes in structural rigidity propagate via three pathways from the cAMP binding sites

The presence of pathways of allosteric signal transmission in proteins has been posited, although the topic is still a matter of debate (33). Here, we used the per-residue free energy Δ*G*_*i*,CNA_ (eq. 4) as a measure of the extent to which a single residue contributes to allosteric signaling (20). The inherent long-range aspect of rigidity percolation makes this analysis an attractive tool for identifying changes in structural stability across a biomolecule’s structure due to distant influences (34, 35).

We perturbed the ensembles of the fully cAMP-bound CL-CNBD by removing all cAMPs (Figure 2A). For ~62% of the residues of the CL-CNBD, Δ*G*_*i,*CNA_ ≥ 0.2 kcal mol^−1^; this threshold was previously applied to detect marked stability changes in this context (20). These residues map out three pathways along which changes in structural stability propagate, starting from the cAMP binding site of each subunit (Figure 2B). The first pathway runs to the adjacent CL-CNBDs along the β-roll of subunit *s* over the β_2_-β_3_ and α_B_-β_7_ loops, and helices A, B, and C in the neighboring subunit *s*+1 (where *s*+*n* with *n* = 0, 1, 2, 3 is the anti-clockwise succession of subunits, seen from the cytosolic site). That way, this pathway mediates the intersubunit signal transmission between neighboring cAMP binding sites. The second pathway also runs to the adjacent CL-CNBDs via the β_2,*s*_ strand of the β-roll to helix D’_*s*+1_ and helix A_*s*+1_; that way, it mediates the intersubunit signal transmission from the cAMP binding site to the “shoulder” motif (36) in the CL region. The third pathway stays within one subunit and proceeds over the strands β_1,*s*_-β_8,*s*_ of the β-roll to helix B’_*s*_ in the CL region close to the TM domain; that way, it mediates the intrasubunit signal transmission from the cAMP binding site to the “elbow” motif in the CL region. Remarkably, residues in helix B’_*s*_ feel the perturbation at the cAMP binding site over a distance of 26 Å.

**Figure 2:**
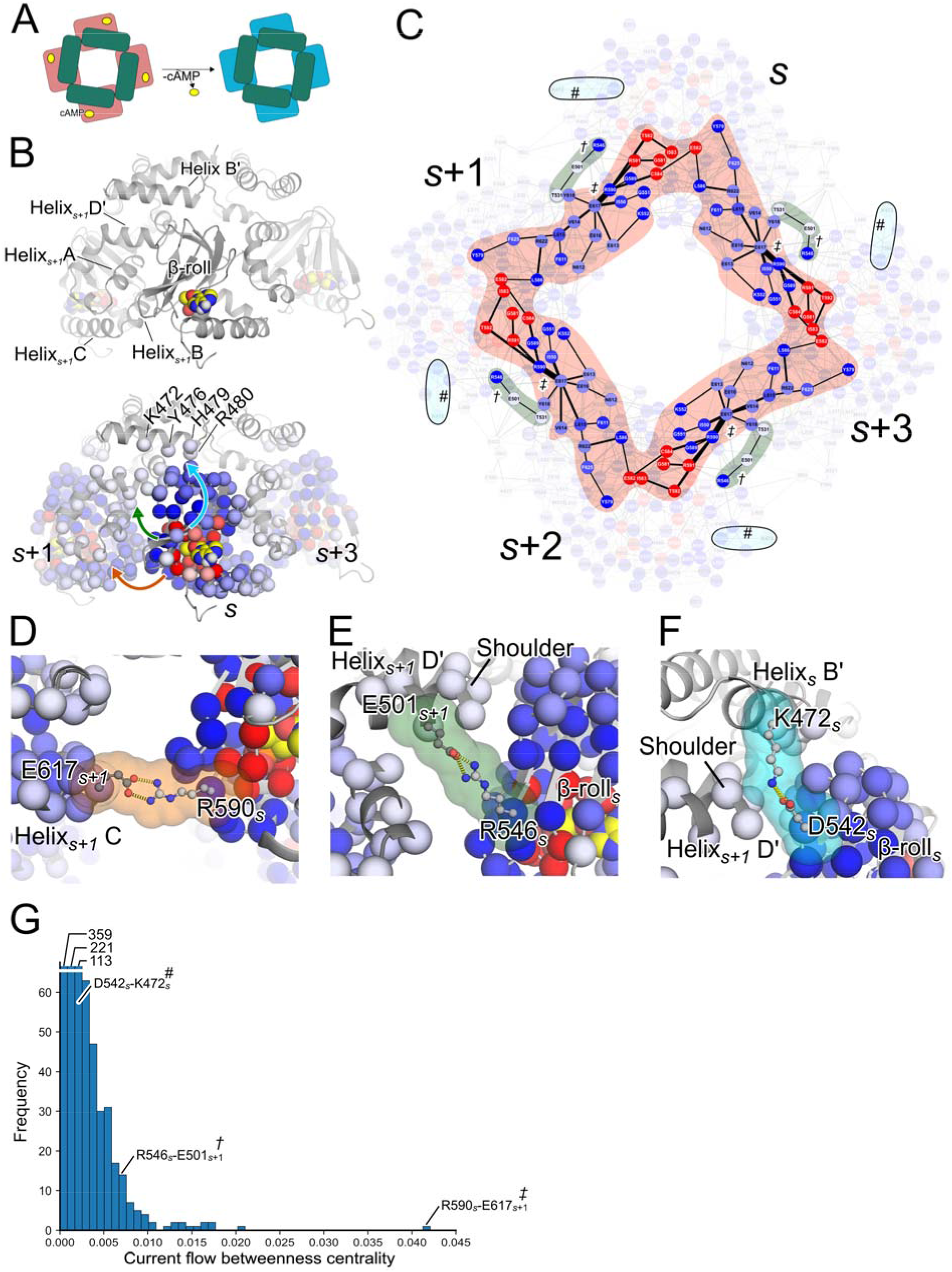
Changes in structural rigidity propagate via three pathways from the cAMP binding sites. (*A*) Perturbation of the CL-CNBD by removal of covalent and non- covalent constraints due to cAMP from the network topology without changing the protein conformation. (*B*) Top: Cartoon representation of the CL-CNBD of mHCN2. Bottom: Residues with significant changes in the structural rigidity due to the removal of cAMP as identified by our perturbation approach are depicted as spheres. Bound cAMP is colored in yellow; residues that are part of the cAMP binding site are colored in red, others in blue. Darker colors indicate a larger change in structural rigidity, Δ*G*_*i,CNA*_. (*C*) The same information from (*B*) is shown as a graph where nodes represent residues and edges the current flow betweenness centrality (eq 4). Highlighted residues are those with the highest centrality values depicting two intersubunit pathways (orange and green). Along both pathways, the two intersubunit salt bridges (*D*) R590_*s*_-E617_*s*+1_ (“*‡*”-label) and (*E*) R546_*s*_-E501_*s*+1_ (“*†*”-label) are located. (*F*) D542_*s*_-K472_*s*_ (“*#*”-label) is part of a third pathway but does not show centrality values that are significantly different from zero. (*G*) The histogram depicts the frequency distribution of centrality values; the first three bars were cut as indicated. Bins are labeled with respect to the three identified salt bridges.

To confirm these dominant routes of signal transmission, we generated a graph consisting of nodes representing the CL-CNBD residues and connecting edges whose weights are defined by the pairwise free energy components Δ*G*_*ij*_ (eq. 4; (20)); the Δ*G*_*ij*_ represent the change in structural stability between pairs of residues upon removal of cAMP(s). We then computed the current flow betweenness centrality *C*_*C*B_(*e*) (37) to identify edges *e* that transmit most of the signal.

Residues with *C*_*C*B_(*e*) values significantly different from zero 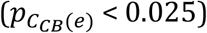 form a ring-like pattern along the β_2,*s*_-β_3,*s*_ and α_B,*s*_-β_7,*s*_ loops and helices A_*s*+1_, B_*s*+1_, and C_*s*+1_, thereby connecting the cAMP binding sites of neighboring subunits (Figure 2C). Along this pathway, the salt bridge between R590_*s*_ and E617_*s* + 1_ connects neighboring subunits (Figure 2D); this salt bridge shows the highest *C*_*C*B_(*e*) value (Figure 2G). R590_*s*_ is located in the β_7_-α_P_ loop, close to the cAMP-binding site, but does not interact with cAMP; E617_*s*+1_ is part of the α_B_-α_C_ loop. The salt bridge persists during the MD simulations of the fully cAMP-bound state with an occurrence frequency of 70.5% (Figure S3A, Table S2).

When considering edges with 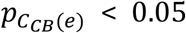 in addition, we observed a second contiguous pattern of three residues between R546_*i*_ in strand β_2,*s*_, E501_*s*+1_ in helix D_*s*+1_, and T531_*s*+1_ in helix A_*s*+1_ (Figure 2B). The residue pair R546_*s*_ and E501_*s*+1_ forms a salt bridge (Figure 2B, E), which persists during the MD simulations (occurrence frequency of 39.1%; Figure S3B, Table S2). It connects the strand β_2,*s*_ with helix D’_*s*+1_ in the CL region. The identified interaction is among the top 4% of those with the highest *C*_*C*B_(*e*) values (Figure 2G).

Along the intrasubunit pathway, changes in structural stability upon removal of cAMP proceed over the strands β_1_-β_8_ within the β-roll. Previous studies demonstrated that β-strands are sensitive to changes in structural stability (20, 38). In comparison to the two intersubunit pathways, edges between residue pairs there do not show *C*_*C*B_(*e*) values that are significantly different from zero (Figure 2B), concordant with a rather broad influence on multiple weakly coupled residues in the β-roll and in contrast to the rather narrow intersubunit pathways. From the β-roll region, the pathway propagates to four residues (K472_*s*_, H479_*s*_, R480_*s*_, and Y476_*s*_) in the CL region in helix B_*s*_’, close to the TM domain of the channel (Figure 2B). K472_*s*_ forms a salt bridge interaction with D542_*s*_ of the β_1,*s*_-β_2,*s*_ loop (Figure 2F), which persists during the MD simulations (occurrence frequency of 66.8%; Figure S3C, Table S2).

To conclude, changes in structural rigidity propagate via three pathways from the cAMP binding sites: On the one hand via a ring-like pattern to neighboring subunits, suggesting that this pathway predominantly influences binding properties of cAMP; on the other hand via a narrow path to the shoulder of the neighboring subunit and via a broad path to the elbow of the same subunit, suggesting that these pathways should predominantly influence cAMP potency because of the involvement of the elbow-on-a- shoulder motif in channel activation (36, 39).

### The intersubunit ring-like pattern impacts cAMP binding affinity and mediates cooperativity between cAMP binding sites but does not directly influence potency

To verify the role of the R590_s_-E617_s+1_ salt bridge within the ring-like pattern, functional studies were carried out. The functional implication of this interaction may be two-fold: First, interrupting it may affect the intersubunit cooperativity and, thus, alter the cAMP binding affinity; second, such an altered cAMP binding may modify the mechanism of cAMP-induced channel gating. We either neutralized (R590A) or reversed (R590E) the positive charge at position 590 to break the salt bridge to E617. We applied confocal patch-clamp fluorometry employing 8-AHT-Cy3B-cAMP (f1cAMP) (Otte et al., 2019) on mHCN2 channel constructs expressed in *Xenopus laevis* oocytes to measure ligand binding (*BC*_50_) (10, 11) and the patch-clamp technique to monitor channel gating (*EC*_50_) (Figure 3, Figure S5).

**Figure 3:**
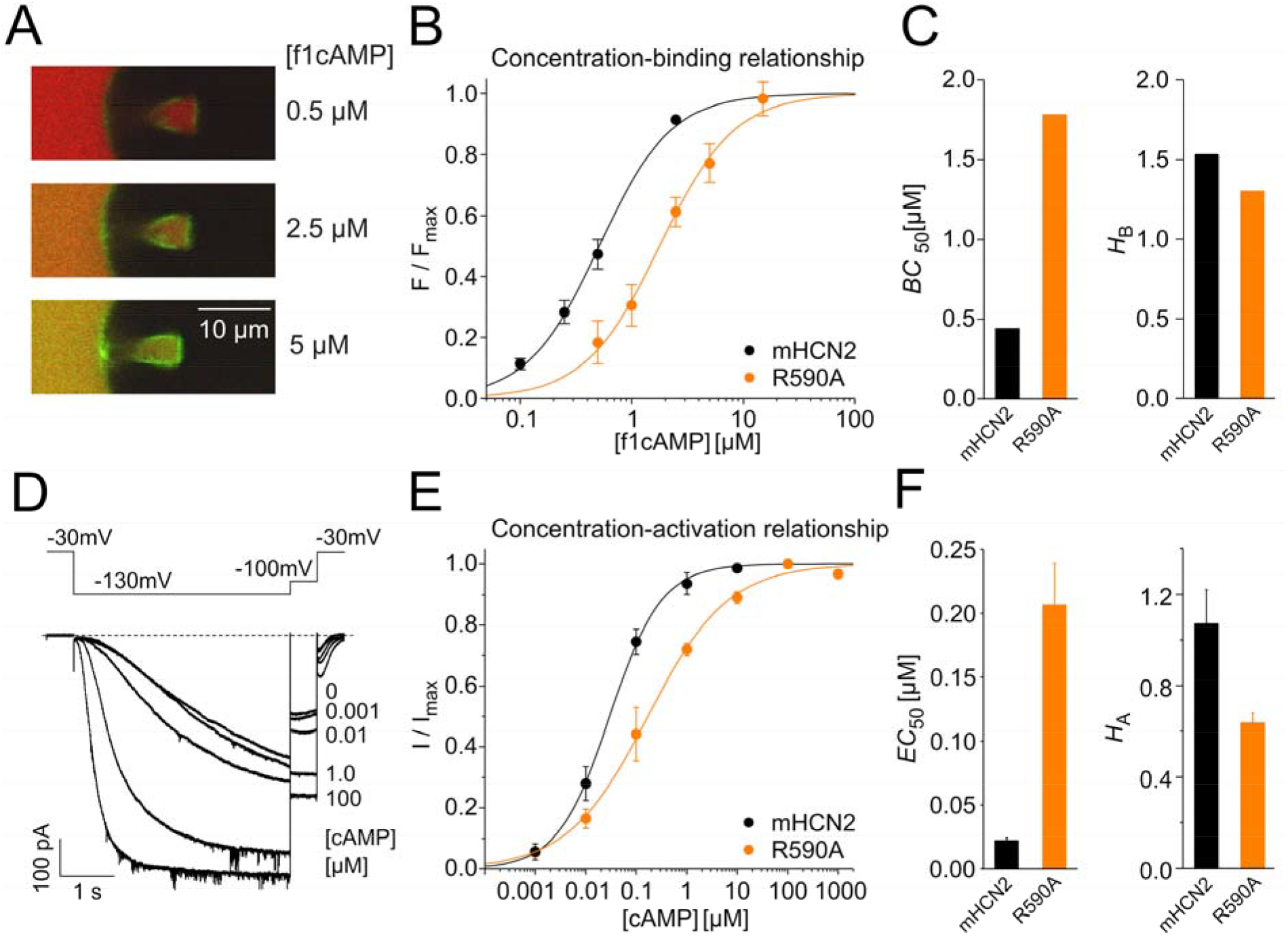
Binding and activation behavior after breaking inter-subunit salt bridges involving R590A. (*A*) Representative confocal images at three different f1cAMP concentrations. Shown is an inside-out patch expressing R590A. The green fluorescence signal arises from the fluorescently tagged ligand f1cAMP, red fluorescence from labeling the background with the reference dye Dy647. (*B*) Concentration-binding relationships obtained from cPCF experiments. Fluorescence intensities *F*, measured at different f1cAMP concentrations at a voltage of -30 mV, were related to the maximum fluorescence signal *F*_max_, measured at a saturating concentration at a voltage of -130 mV. *F*/*F*_max_ values are averages from 3 to 6 recordings. Solid lines represent the results of fitting the Hill equation (eq. 8) to the data, yielding the concentration of half-maximum binding, *BC*_50_, and the Hill coefficient of binding, *H*_B_. (*C*) Left panel: *BC*_50_ values for mHCN2 and R590A. Right panel: *H*_B_ values for mHCN2 and R590A. (*D*) Representative traces at different cAMP concentrations from R590A. The voltage protocol is shown above. (*E*) Concentration-activation relationships from excised inside-out macropatches. Current amplitudes *I* were related to the maximum current amplitude *I*_max_ at a saturating concentration in each patch. The Hill equation was approximated to the data of each recording, yielding the concentration of half-maximum activation, *EC*_50_, and the Hill coefficient of activation, *H*_A_. (*F*) Left panel: *EC*_50_ values for mHCN2 and R590A. Right panel: *H*_A_ values for mHCN2 and R590A. Error bars indicate SEM. All fitted parameters regarding binding and activation are summarized in Table S3.

First, we tested specific ligand binding in R590A (Figure 3A-C) at concentrations between 0.1 μM and 5 μM f1cAMP in inside-out macropatches via the bath solution at a non-activating voltage of -30 mV (Figure 3A; Material and Methods, (10)). The channel-bound fraction was then normalized to the maximal intensity at 2.5 μM for mHCN2 and 15 μM for R590A, both at -130 mV, and plotted versus the respective concentration (Figure 3B). The Hill equation (eq. 8) was fitted to the averaged data, yielding the concentration of half-maximum binding, *BC*_50_, and the Hill coefficient, *H*_B_ (Figure 3B, C). The binding affinity was ~4-fold lower in R590A compared to mHCN2 wildtype. The Hill coefficient decreased slightly by ~15%. A similar result was obtained when replacing the respective arginine by glutamate (Figure S4). All fit results are summarized in Table S3. Second, we tested the effect on channel activation in inside-out macropatches at cAMP concentrations ranging from 1 nM to 1000 μM with the voltage protocol in Figure 3D.

Normalized tail currents were plotted versus the cAMP concentration, and the Hill equation (eq. 7) was fitted to the data points of each recording. The concentrations of half-maximum activation, *EC*_50_, and Hill coefficients of activation, *H*_A_, were determined (Figure 3E, F). For R590A, the *EC*_50_ value was ~9-fold higher than for mHCN2, indicating a reduced potency. This higher *EC*_50_ value can arise from the impaired binding shown in Figure 3B, C. *H*_A_ in R590A was significantly reduced compared to mHCN2, which suggestsa reduced cooperativity in gating and, eventually, in ligand binding. A similar result was found when replacing the respective arginine by glutamate, with the exception that *H*_A_ in R590E was less reduced than in R590A (Figure S4). All fit results are summarized in Table S3.

To conclude, the intersubunit ring-like pattern involving the R590_*s*_-E617_*s*+1_ salt bridge impacts cAMP binding affinity, mediates cooperativity between cAMP binding sites, and, thereby, indirectly influences potency.

### The intersubunit pathway to the shoulder and the intrasubunit pathway to the elbow influence potency but neither cAMP binding affinity nor cooperativity between cAMP binding sites

To verify the role of the R546_*s*_-E501_s+1_ salt bridge within the pathway to the shoulder, functional studies were carried out in R546A and R546E. In analogy to the experiments for R590A and R590E, binding and gating were studied by cPCF and patch-clamp (Figure 4A-D). *BC*_50_ for R546A was increased while *H*_B_ was similar to mHCN2 (Figure 4A, B), suggesting that the binding affinity was reduced but the cooperativity was not affected. Surprisingly, *EC*_50_ for R546A was significantly lower (~8- fold) than for mHCN2, indicating a higher potency in this construct (Figure 4C, D). Because cAMP binding to R546A is hampered, this higher potency must be caused by improved gating rather than by stronger binding. A similar result, but with a less pronounced increase in binding affinity, was found when replacing the respective arginine by glutamate (Figure S4).

**Figure 4:**
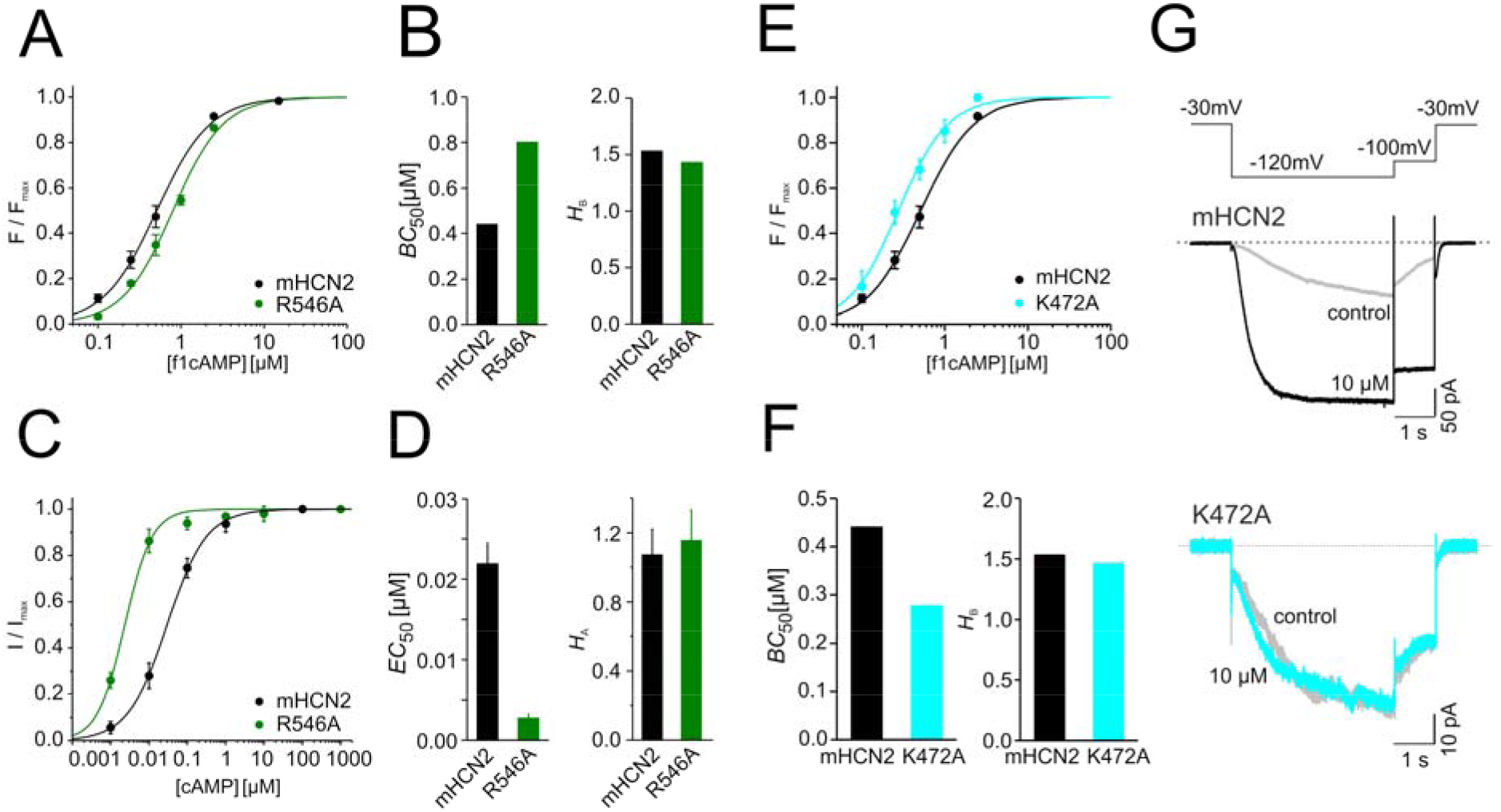
Binding and activation after neutralizing R546 and K472. (*A*) Concentration-binding relationship obtained from cPCF experiments for R546A. Fluorescence intensities, *F*, measured at different f1cAMP concentrations at -30 mV, were related to the maximum fluorescence signal *F*_max_, measured at a saturating concentration of 2.5 μM f1cAMP at a saturating voltage of -130 mV. *F*/*F*_max_ values are averages from 3 to 6 recordings. Solid lines represent the result of approximating the Hill equation (eq. 8) to the data, yielding the concentration of half-maximum binding, *BC*_50_, and the Hill coefficient of binding, *H*_B_. Error bars indicate SEM. (*B*) Left panel: *BC*_50_ values for mHCN2 and R546A. Right panel: *H*_B_ values for mHCN2 and R546A. (*C*) Concentration-activation relationships from inside-out macropatches. Current amplitudes *I* were related to the maximum current amplitude *I*_max_ at a saturating concentration in each patch. The Hill equation was approximated to the data of each recording, yielding the concentration of half-maximum activation, *EC*_50_, and the Hill coefficient of activation, *H*_A_. (*D*) Left panel: *EC*_50_ values for mHCN2 and R546A. Right panel: *H*_A_ values for mHCN2 and R546A. (*E*) Concentration-binding relationship obtained from cPCF experiments for K472A in analogy to panel A. (*F*) Left panel: *BC*_50_ values for mHCN2 and K472A. Right panel: *H*_B_ values for mHCN2 and K472A. (*G*) Representative current traces for mHCN2 (upper panel) and K472A (lower panel) in the absence (control) and presence of 10 μM cAMP. The voltage pulse protocol is indicated above. All fitted parameters regarding binding and activation are summarized in Table S3.

To study the role of the salt bridge formed by K472 with D542 within the intrasubunit pathway, we replaced lysine by alanine for neutralizing the charge. First, we quantified ligand binding using cPCF (Figure 4E, F; Table S3), in a similar way as described for R546 and R590 mutants (Figure 4A, B). The binding affinity, *BC*_50_, was higher by ~1.9- fold than for mHCN2, while the Hill coefficient, *H*_B_, for binding was not changed, indicating no effect on cooperativity between cAMP binding sites. Regarding cAMP-induced activation, we observed that K472A is not responsive anymore, i.e., neither the current amplitude nor the activation kinetics was changed (Figure 4G).

To conclude, the intersubunit pathway to the shoulder motif involving the R546_*s*_- E501_*s*+1_ salt bridge influences the potency, but impacts the cAMP binding affinity to a much lesser extent than R590_*s*_-E617_*s*+1_. Hence, this pathway is not involved in mediating cooperativity between cAMP binding sites. The effects of distorting the intrasubunit pathway are similar to those observed when distorting the intersubunit pathway to the elbow motif in that potency is influenced, the cAMP binding affinity increases weakly, and the cooperativity between cAMP binding sites remains unchanged. Yet, while distorting the intersubunit pattern causes a higher potency in this construct, distorting the intrasubunit pathway abrogates cAMP potency. According to ref. (12), the latter result suggests that the mutant shifts the channel from the low-affinity to the high-affinity state, which would also explain the increase in the cAMP binding affinity.

## Discussion

In this study, we addressed the question how changes in the structural dynamics of the CL-CNBD of mHCN2 upon cAMP binding relate to inter- and intrasubunit allosteric signal transmission. We combined extensive MD simulations in combination with a rigidity theory-based free energy perturbation approach to scrutinize the role of dynamic, i.e., entropically dominated, allostery for cooperativity within and between the CL-CNBDs of the subunits, study signal transmission evoked by bound cAMP, and identify interactions involved in the signaling. The rigidity theory-based free energy perturbation approach (20) proved successful in related studies of membrane proteins (40-42). Our results allowed us to map out distinct routes within the tetrameric CL-CNBD that modulate different cAMP binding responses in HCN2 channels through disjunct salt bridge interactions.

First, our computational results yielded for the binding of two, three, and four cAMP molecules to tetrameric CL-CNBD in the non-activated conformation the succession of negative, no, and positive for the cooperativity. These results are in perfect semi-quantitative agreement with results from fitting Markovian models to experimental time courses of cAMP-binding and unbinding (13) and are also concordant with previous results that stressed the complex nature of cooperativity in HCN channels (13, 43, 44). Yet, whereas the experiments (13, 29, 43, 44) used full-length mHCN2 channels, here we analyzed only the tetrameric CL-CNBD. Hence, our results demonstrate that non-uniform changes in structural rigidity of only the CL-CNBD are sufficient to give rise to the complex succession of cooperativities for cAMP binding steps. Our results vary, however, from others showing a different succession of cooperativities in a truncated CL-CNBD fragment (45).

Our computational results, which originate from rigidity percolation upon cAMP binding, are also concordant with tmFRET and DEER experiments (14-17) that showed a narrowing of distance distributions upon cAMP binding between residues in helix C and the β-roll in the CNBD, indicative of a decreased number of accessible configurational states of the CL-CNBD. They furthermore provide an explanation at the atomistic level for why the structurally equal CL-CNBD subunits are non-equal in their function (46) and show that states with two and four cAMP are structurally more stable than those with one and three cAMP molecules, respectively (Figure 1D). While this is in line with a dimer-of-dimers organization, as suggested previously (46), both the finding that ΔΔ*G*_2_ is equal for *cis* and *trans* configurations with two bound cAMP molecules, and that the third binding event is associated with no cooperativity, indicate that in non-activated tetrameric CL-CNBD the dimers only cooperate when fully ligated. This result reconciles earlier ones that either found no cooperativity between dimers (29, 47) or cooperativity for three and four cAMP bound in the case of activated HCN channels (46).

Second, our computational results revealed changes in structural rigidity that propagate via three pathways from the cAMP binding sites. Cooperativity in biomolecules usually relies on the long-range coupling between distal functional elements (48). Such a long-range coupling can be achieved by a small set of salt bridges that mediate the equilibrium between ligand binding and tertiary and quaternary structural dynamics (46). Indeed, rigorous graph-based analyses revealed for the ring-like pattern to neighboring subunits and the one that propagates via a narrow path to the shoulder of the neighboring subunit one salt bridge each (R590_*s*_ and E617_*s*+1_; R546_*s*_ and E501_*s*+1_) through which most of the information flows. On the broad path to the elbow of the same subunit, another salt bridge (K472_*s*_ and D542_*s*_) was identified. The existence of the pathways is concordant with previous studies that demonstrated pronounced cooperativity among the four CL-CNBD subunits (13, 44) as well as allosteric feedback from the TM channel core to the cAMP binding sites in the CL-CNBD (49). Yet, the exact course of the allosteric signal transmission remained elusive in those studies.

Electrophysiological and patch-clamp fluorometry experiments showed that each of the three salt bridges alters the gating of mHCN2 channels if the interaction is lost. Notably, mutational analyses showed that depending on which salt bridge is broken, different changes in the mechanism of channel activation are observed. These findings strengthen the idea of the modular structure of HCN channels where each module takes over a specific function (50). They also support our predictions of the functional influence of these pathways. As to the ring-like pattern, the loss of the R590_*s*_-E617_*s*+1_ salt bridge led to an indirectly reduced potency compared to mHCN2 wildtype channels via a reduced cAMP affinity. By contrast, loss of the second intersubunit salt bridge between R546_*s*_ and E501_*s*+1_ led to a higher potency in this variant. Since the channel’s affinity for cAMP was reduced compared to the wildtype channel, the higher potency must be caused by improved gating rather than by enhanced cAMP binding. Finally, loss of the intrasubunit salt bridge between K472_*s*_-D542_*s*_ led to an increased affinity of the channel for cAMP in conjunction with abrogating the cAMP potency. Surprisingly, the loss of one of the three salt bridges did not change the Hill coefficients for binding markedly. Yet, applying a model for cooperative binding to four binding sites (see Supplementary Text) shows that homogeneous or heterogeneous changes in microscopic association constants on the order of the *BC*_50_ changes observed herein (1.9-fold to 4.0-fold) do not lead to marked changes of the Hill coefficient either (Figure S6).

Early studies of the HCN CL-CNBD already aimed at identifying amino acids crucial for regulation of cAMP-dependent channel gating (Table S4). Most of those amino acids are within the cNMP binding site. Additionally, four interactions distant to the cNMP binding site have been reported: The two interactions R590_*s*_-E617_*s* + 1_ and K472_*s*_-D542_*s*_ identified here have been reported previously to be relevant for channel gating (51, 52) and, thus, substantiate our computations. By contrast, the interactions K452_*s*_-E494_*s*+1_ and K472_*s*_- E502_*s*+1_, identified previously by mutational studies (51, 52), were not identified by our computations. E494 is located on helix A’ such that its functional role may be predominant in the activated channel, which we have not investigated here. This is supported by a low occurrence frequency of the K452_*s*_-E494_*s*+1_ interaction during our MD simulations (Figure S3D). Likewise, the occurrence frequency of the K472_*s*_-E502_*s*+1_ interaction is low, as, in turn, the competing K472_*s*_-D542_*s*_ interaction prevails (Figure S3C, E). The R546_*s*_-E501_*s*+1_ interaction has been investigated in detail in this study for the first time.

In summary, we mapped out distinct routes within the CL-CNBD that modulate different cAMP binding responses in HCN2 channels through disjunct salt bridge interactions. With respect to other studies investigating the role of functionally relevant modules in HCN channel function (17, 53-56), our results signify that functionally relevant submodules may exist within and across structurally discernable subunits.

## Materials and methods

### Molecular dynamics simulations

#### Tetrameric CL-CNBD from the mouse HCN2 channel

The initial structure was taken from PDB id 1q5o, and selenomethionine residues were replaced by methionine residues. From this fully cAMP-bound structure, we derived the configurations with no, one, two (cis/trans), and three cAMP molecules bound. All these structures served as starting structures for all-atom MD simulations of 1 μs length, performed in 10 replica per system, resulting in, in total, 60 μs of simulation time. Details on the setup and execution of MD simulations of the isolated tetrameric CL-CNBD are given in the Supplementary Methods.

### Constraint network analysis

Rigidity analyses were performed using the CNA software package (21). We applied CNA on ensembles of network topologies generated from conformational ensembles provided as input (see Supplementary Methods). Average stability characteristics are calculated by constraint counting on each topology in the ensemble. The FIRST software (version 6.2) (57), to which CNA is a front and back end, was used to construct each network of covalent and non-covalent (hydrogen bonds including salt bridges and hydrophobic tethers) constraints. The hydrogen bond energy *E*_HB_ determined from an empirical energy function (58) describes the strength of hydrogen bonds (including salt bridges). Hydrophobic tethers between carbon and sulfur atoms were taken into account if the distance between these atoms was less than the sum of their van der Waals radii (C: 1.7 Å, S: 1.8 Å) plus *D*_*cut*_ = 0.25 Å (26).

Biomolecules generally display a hierarchy of structural stability, reflecting the modularity of their structure (22). To identify this hierarchy, a “constraint dilution trajectory” of network states {*σ*} is analyzed, obtained from an initial network topology by successively removing non-covalent constraints (22-26). Here, hydrogen bond constraints (including salt bridges) are removed in the order of increasing strength such that for network state *σ,* only those hydrogen bonds are retained that have an energy *E*_*HB*_ ≤ *E*_*cut*_(*σ*).

### Allosteric signaling through constraint networks

We used a per-residue decomposition scheme to identify the extent to which single residues contribute to the allosteric signaling. First, neighbor stability maps (*rc*_*ij*,neighbor_) are derived from the “constraint dilution trajectory”; they contain information accumulated over all network states {σ} along the trajectory (20, 59). More precisely, stability maps monitor the persistence of rigid contacts for pairs of residues during a bond dilution process; a rigid contact is present as long as two residues belong to the same rigid cluster *c* of the set of rigid clusters C^*Ecut*^. As our focus is on short-range rigid contacts, only pairs of heavy atoms of the residue pair *R*_{*i, j*}_, *A*_{*k* ∈ *i*, *l* ∈ *j*}_, separated by a distance *d* ≤ 4.5 Å are considered. The resulting neighbor stability map (eq 2) (59) relates to the local stability in the network.

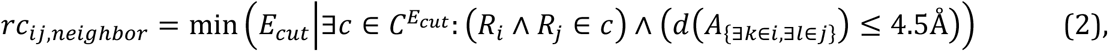

A chemical potential energy *E*_*i*,CNA_ of residue *i* is obtained by summation over all *n* short-range rigid contacts in which the residue is involved (eq 3)

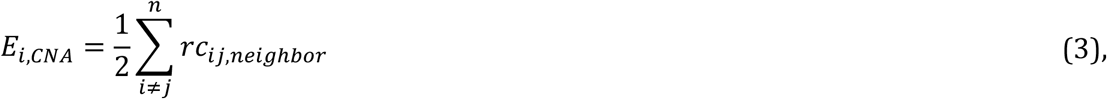

which is then used in a linear response approximation to obtain the per-residue decomposition (eq 4)

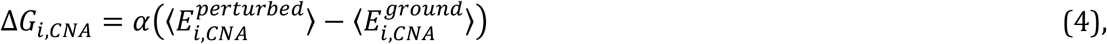

with α being ½ (20).

For identification of the most dominant signaling pathways through the isolated tetrameric CL-CNBD, we computed the current flow betweenness centrality to quantify the relative importance of each site concerning the information flow in a graph. Stability maps as derived from rigidity analysis were used to generate an undirected graph *G* = (*V*, *E*). In such a graph, nodes *V* represent the residues and edges *E* the pairwise perturbation free energy *ΔG*_*ij*_ (eq. 5) between residue *i* and *j* on going from a cAMP-bound to an *apo* state

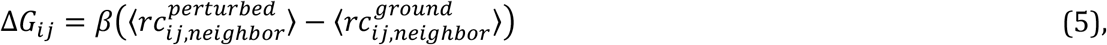

where 〈*rc*_*ij, neighbor*_ 〉 is the average strength of a rigid contact (eq 2) and β= ½ as used in ref (20). The current flow betweenness centrality *C*_*C*B_(*e*) of each edge *e*, starting from a source *s* ∈ *V* to a sink *t* ∈ *V*, is then computed for all combinations of starting point *s* and target *t* from the throughputs 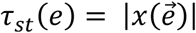 due to a *st*-current (eq 6) (37)

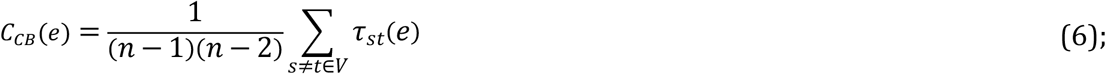

n = ∥ V ∥, and the term in front of the summation sign is a normalizing constant with respect to the size of the graph. For the graph construction and calculation of the current flow betweenness centrality, the NetworkX extension for Python was used. The underlying algorithm to calculate the current flow betweenness centrality is described in ref (60).

### Oocyte preparation

Oocytes were surgically removed from adult females of the South African clawed frog *Xenopus laevis* under anesthesia (0.3% MS-222 (tricaine methanesulfonate) (Pharmaq Ltd. Fordingbridge, UK)). The oocytes were treated with collagenase A (3 mg/ml; Roche (Grenzach-Wyhlen, Germany)) for 105 min in Ca^2+^-free Barth’s solution containing (in mM) 82.5 NaCl, 2 KCl, 1 MgCl_2_, and 5 HEPES, pH 7.5. Oocytes of stages IV and V were manually dissected and injected with cRNA encoding mHCN2 channels of Mus musculus or constructs carrying a point mutation. After injection with cRNA, the oocytes were cultured at 18°C for 2–10 d in Barth’s solution containing (in mM) 84 NaCl, 1 KCl, 2.4 NaHCO_3_, 0.82 MgSO_4_, 0.41 CaCl_2_, 0.33 Ca(NO_3_)_2_, 7.5 Tris, cefuroxime, and penicillin/streptomycin, pH 7.4. The procedures had approval from the authorized animal ethics committee of the Friedrich Schiller University Jena. The methods were performed by the approved guidelines. Oocytes harvested in our laboratory were complemented with ready-to-use oocytes purchased from EcoCyte Bioscience (Dortmund, Germany).

### Molecular biology

The mouse HCN2 (UniProt ID O88703 including two modifications, G8E and E55G without functional relevance) channel and all modified subunit variants were subcloned in front of the T7 promoter of pGEMHEnew. Point mutations (K472A, R546A, R546E, R590A, R590E) were introduced via the overlapping-PCR strategy followed by subcloning of the modified fragment using flanking restriction sites. The correctness of the sequences was confirmed by restriction analysis and sequencing (Microsynth SEQLAB, Göttingen, Germany). cRNAs were prepared using the mMESSAGE mMACHINE T7 Kit (Ambion Inc, Austin, USA).

### Electrophysiology

Ionic currents were measured with the patch-clamp technique in inside-out macropatches. The patch pipettes were pulled from quartz tubing (P-2000; Sutter Instrument, Novato, US) whose outer and inner diameters were 1.0 and 0.7 mm, respectively (Vitrocom, Silsden, UK). The pipette resistance was 0.9-2.7 MΩ. The bath solution contained (in mM) 100 KCl, 10 EGTA, and 10 HEPES, pH 7.2. The pipette solution contained (in mM) 120 KCl, 10 HEPES, and 1.0 CaCl_2_, pH 7.2. For measuring concentration-activation relationships, different cAMP concentrations (BIOLOG LSI GmbH & Co KG, Bremen, Germany) were added to the bath solution.

An HEKA EPC 10 USB amplifier (Harvard Apparatus, Holliston, US) was used for the current recording. Pulsing and data recording were controlled by the Patchmaster software (Harvard Apparatus, Holliston, US). The sampling rate was 5 kHz. The holding potential was generally -30 mV. Each recording was started 3–4 min after patch excision to avoid artifacts caused by excision-induced channel rundown (13, 61).

### Confocal patch-clamp fluorometry

The fluorescence intensity in the patch was measured by patch-clamp fluorometry (62, 63) combined with confocal microscopy (10, 12). As a fluorescent ligand, we used 8-AHT- Cy3B-cAMP (f1cAMP), a cAMP to which the fluorescent dye Cy3B (GE Healthcare, Frankfurt, Germany) was linked via an aminohexylthio spacer to position 8 of the adenosine moiety (64, 65). The recordings were performed with an LSM 710 confocal microscope (Carl Zeiss AG, Jena, Germany) and were triggered by the ISO3 software (MFK, Niedernhausen, Germany). To distinguish the fluorescence of the non-bound f1cAMP from that of the bound f1cAMP, a second, chemically related dye, DY647 (Dyomics GmbH, Jena, Germany), was also added to the bath solution. The 543 nm and 633 nm lines of a He-Ne laser were used to excite f1cAMP and DY647, respectively. For quantifying the bound fcAMP, the fluorescence intensities of the red and the green channels were corrected for small offsets, and the fluorescence in the red channel was scaled to the fluorescence in the green channel in the bath. The difference between the measured green and the scaled red profile for each pixel of the confocal image represents the fraction of the fluorescence signal originating from the bound f1cAMP. The free patch membrane (patch dome) only was used to quantify binding by setting a mask at a region of interest. The fluorescence, *F*, was averaged over all pixels inside this mask and normalized in each patch concerning the fluorescence at saturating [f1cAMP] and full channel activation (−130 mV), *F*_max_. The recording rate of the confocal images was 10 Hz.

### Data analysis

Concentration-activation relationships were analyzed by fitting the Hill equation (eq 7) to the individual current data using the OriginPro 9.0G software (Northampton, USA):

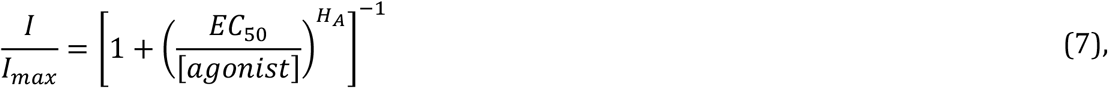

with *I* being the actual current amplitude, *I*_max_ the maximal current amplitude at a saturating ligand concentration, *EC*_50_ the concentration of half-maximum activation, and *H*_A_ the Hill coefficient of activation. Experimental data are given as mean ± SEM.

Concentration-binding relationships were analyzed by fitting the Hill equation (eq 8) to the mean fluorescence data using the OriginPro 9.0G software (Northampton, USA):

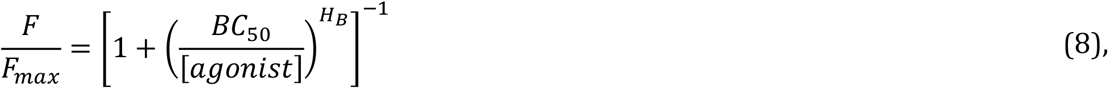

with *F* being the actual fluorescent intensity, *F*_max_ the maximal fluorescent intensity at a saturating ligand concentration and saturating voltage, *BC*_50_ the concentration of half-maximum binding, and *H*_B_ the Hill coefficient of binding.

### Statistical analysis of computational results

Statistical analyses were performed with the Python modules numpy 1.16.4 and scipy 1.3.0. The data is given as mean ± SEM. For ΔΔ*G*_*CNA,n*_ values, SEM were computed following the law of error propagation for the addition of measured quantities according to eq. 8

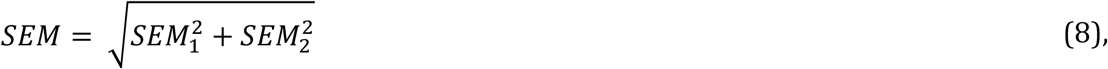

where *SEM*_{1,2}_ are the individual errors of two measures, respectively.

The two-sided Welch’s *t*-test as implemented in scipy was used to evaluate the significance of differences between mean values.

### Statistical analysis of experimental results

Experimental data are given as mean ± SEM.

## Acknowledgments

This work was supported by the grant Research Unit 2518 DynIon of the Deutsche Forschungsgemeinschaft (DFG) (projects P2 (KU 3092/2-2; BE1250/19-1) and P7 (GO 1367/2-2)). We thank Sandra Bernhardt, Uta Enke, Andrea Kolchmeier, Claudia Ranke and Karin Schoknecht for excellent technical assistance. We are grateful for computational support and infrastructure provided by the “Zentrum für Informations- und Medientechnologie” (ZIM) at the Heinrich Heine University Düsseldorf and the computing time provided by the John von Neumann Institute for Computing (NIC) to HG on the supercomputer JUWELS at Jülich Supercomputing Centre (JSC) (user IDs: HKF7; HDD17).

