## Supplemental Information for "Pathways of inter- and intrasubunit allosteric signaling in the C-linker disk and cyclic nucleotide-binding domain of HCN2 channels"

**This PDF file includes:**

Supplementary text
Figures S1 to S6
Tables S1 to S4
SI References

**Supplementary Text**

**Hill coefficient for *N* = 4 binding sites.** We applied a systematic treatment of site-specific effects in systems composed of a small number of binding sites. According to ref. (1), a system with $N = 4$ binding sites is the simplest case that shows the role of second-order contracted partition functions in the resolution of site-specific parameters. From this, we computed the Hill coefficient $H_B$ for mHCN2 to quantify how strongly changes in intersubunit cooperativity and, hence, changes in microscopic binding affinities of cAMP affect $H_B$.

$$H_B = 1 + \frac{3K_{A2}\,[L]\left(1+2K_{A3}[L]+K_{A3}K_{A4}[L]^2\right)}{1+3K_{A2}[L] + 3K_{A2}K_{A3}[L]^2 + K_{A2}K_{A3}K_{A4}[L]^3} - \frac{3K_{A1}\,[L]\left(1+2K_{A2}[L]+K_{A2}K_{A3}[L]^2\right)}{1+3K_{A1}[L] + 3K_{A1}K_{A2}[L]^2 + K_{A1}K_{A2}K_{A3}[L]^3} \qquad \text{eq. S1}$$

Here, [L] is the ligand concentration in M, and $K_{Ax}$ with $x$ = 1, 2, 3, 4 are equilibrium association constants for the respective binding step $x$ of cAMP to inactivated mHCN2 taken from ref. (2):

$K_{A1} = 4.94 \times 10^5\ M^{-1}$
$K_{A2} = 1.69 \times 10^5\ M^{-1}$
$K_{A3} = 1.69 \times 10^5\ M^{-1}$
$K_{A4} = 2.91 \times 10^6\ M^{-1}$

Fig. S6 depicts how changing the $K_{AX}$ impacts the cooperativity and overall Hill coefficient $H_B$ in mHCN2.

### Supplementary Methods

#### Molecular dynamics simulations

All MD simulations of the isolated tetrameric CL-CNBDs were carried out with the AMBER 17 package of molecular simulation programs (3) using the GPU-accelerated version of PMEMD (4). The ff14SB force field for proteins (5) and the GAFF force field for ligands (6) were employed. The structures were solvated in a truncated octahedron of TIP3P water (7) such that the distance between the boundary of the box and the closest solute atom was at least 11 Å. Periodic boundary conditions were applied using the particle mesh Ewald (PME) method (8) to treat long-range electrostatic interactions. Bond lengths involving bonds to hydrogen atoms were constrained by SHAKE (9). The direct-space non-bonded cutoff was 8 Å. In the equilibration phase, first, the solvent was minimized for 250 steps by using the steepest descent method followed by conjugate gradient minimization of 50 steps. Subsequently, the same approach was used to minimize the entire system, including the protein. Afterwards, the system was heated from 100 K to 300 K using canonical ensemble (NVT) MD simulations, and the solvent density was adjusted using isothermal-isobaric ensemble (NPT) MD simulations. Positional restraints applied during thermalization were reduced in a stepwise manner over 50 ps, followed by 50 ps of unrestrained isothermal-isobaric ensemble (NPT) MD simulations at 300 K with a time constant of 1 ps for heat bath coupling. The time step was set to 2 fs. The backbone atoms of the residues D443-R447 of each subunit were kept positionally restrained throughout the equilibration and production phase to mimic the presence of a TM domain. Each production phase was carried out using an increased time step of 4 fs with hydrogen mass repartitioning (10). The simulation time was 1 µs using 1 ps for heat bath coupling, and coordinates were saved at 500 ps intervals resulting in ensembles of 2000 conformations.

**Supplementary Figures**

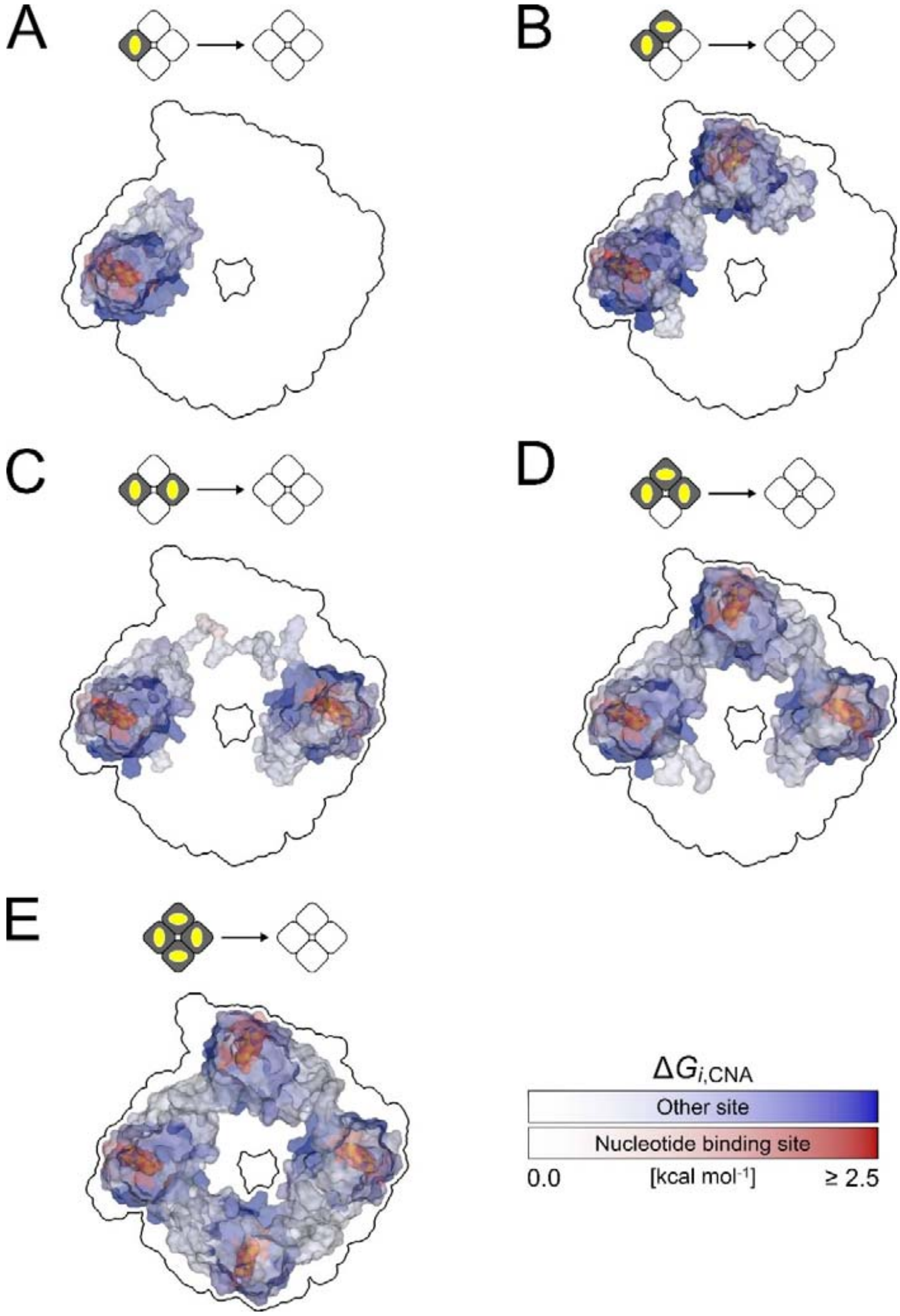


**Fig. S1.** Per-residue free energies $\Delta G_{i,CNA}$ (eq. 5) depicting changes in biomolecular stability of the tetrameric CL-CNBD upon removal of (A) 1, (B) 2 (cis), (C) 2 (trans), (D) 3, and (E) 4 cAMP from the corresponding ligand-bound states as perturbation, mapped onto the structure of the tetrameric CL-CNBD viewed from the cytosolic site. The color indicates if a residue is part of the nucleotide binding site (red) or any other site (blue). Darker colors indicate larger $\Delta G_{i,CNA}$ values (see color scale). The black outline depicts the contour of the CL-CNBD.

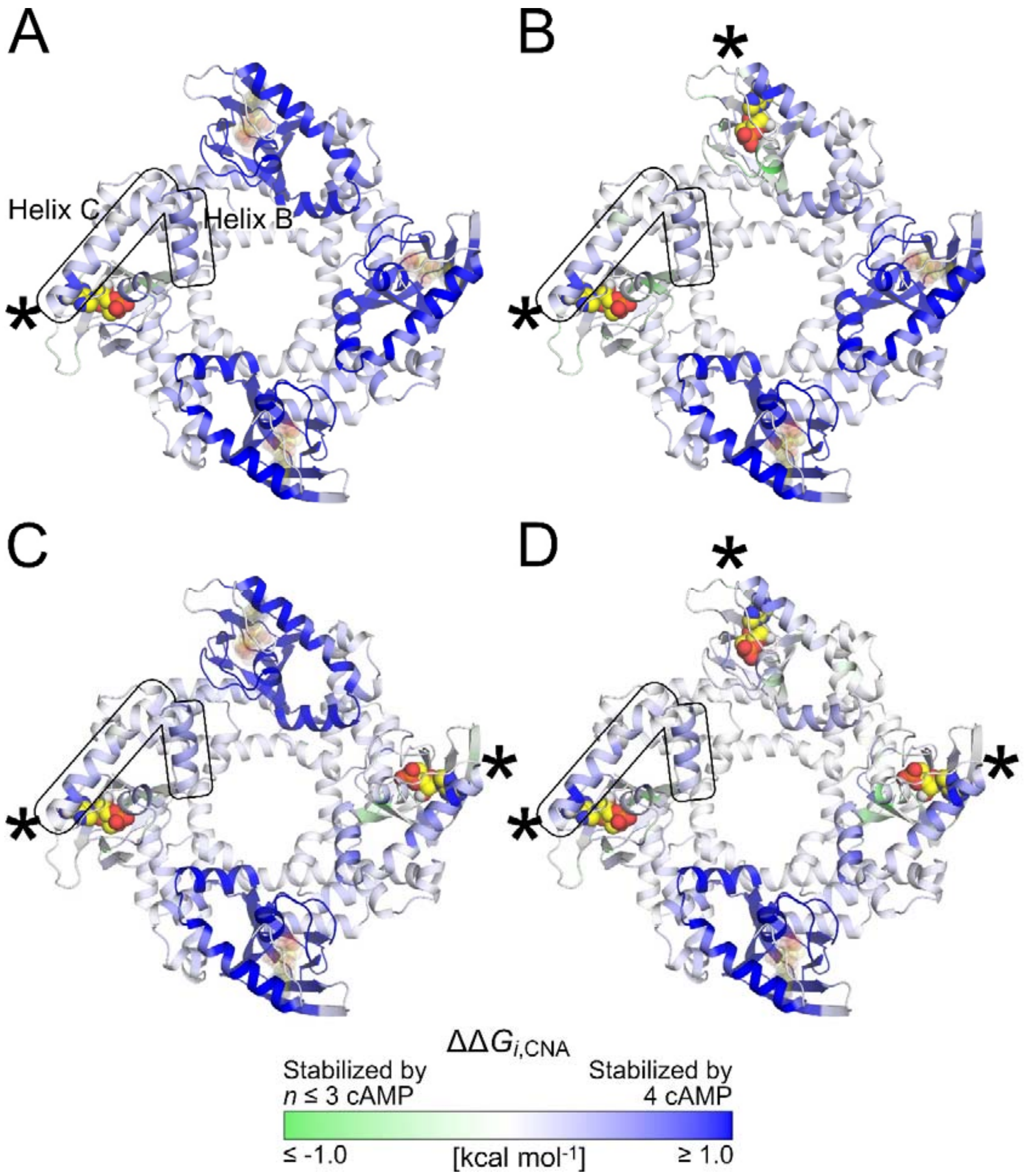


**Fig. S2.** Differences between per-residues free energies $\Delta G_{i,\mathrm{CNA}}$ for the single (A), double (cis (B) / trans (C)), and triple (D) cAMP-bound state *versus* the fully liganded state, $\Delta\Delta G_{i,\mathrm{CNA}}$, mapped onto the tetrameric CL-CNBD structure viewed from the cytosolic site. Bluish colors indicate regions of the CL-CNBD that are more stabilized by the fully liganded state than one of the other states; greenish colors indicate the opposite. Asterisks label the subunits where the cAMP is present in the single, double, and triple bound states. The results show that the fully liganded state leads to an extra stabilization particularly of helices B and C (highlighted for one subunit using a black outline) even in subuits to which cAMP was bound already (marked by asterisks).

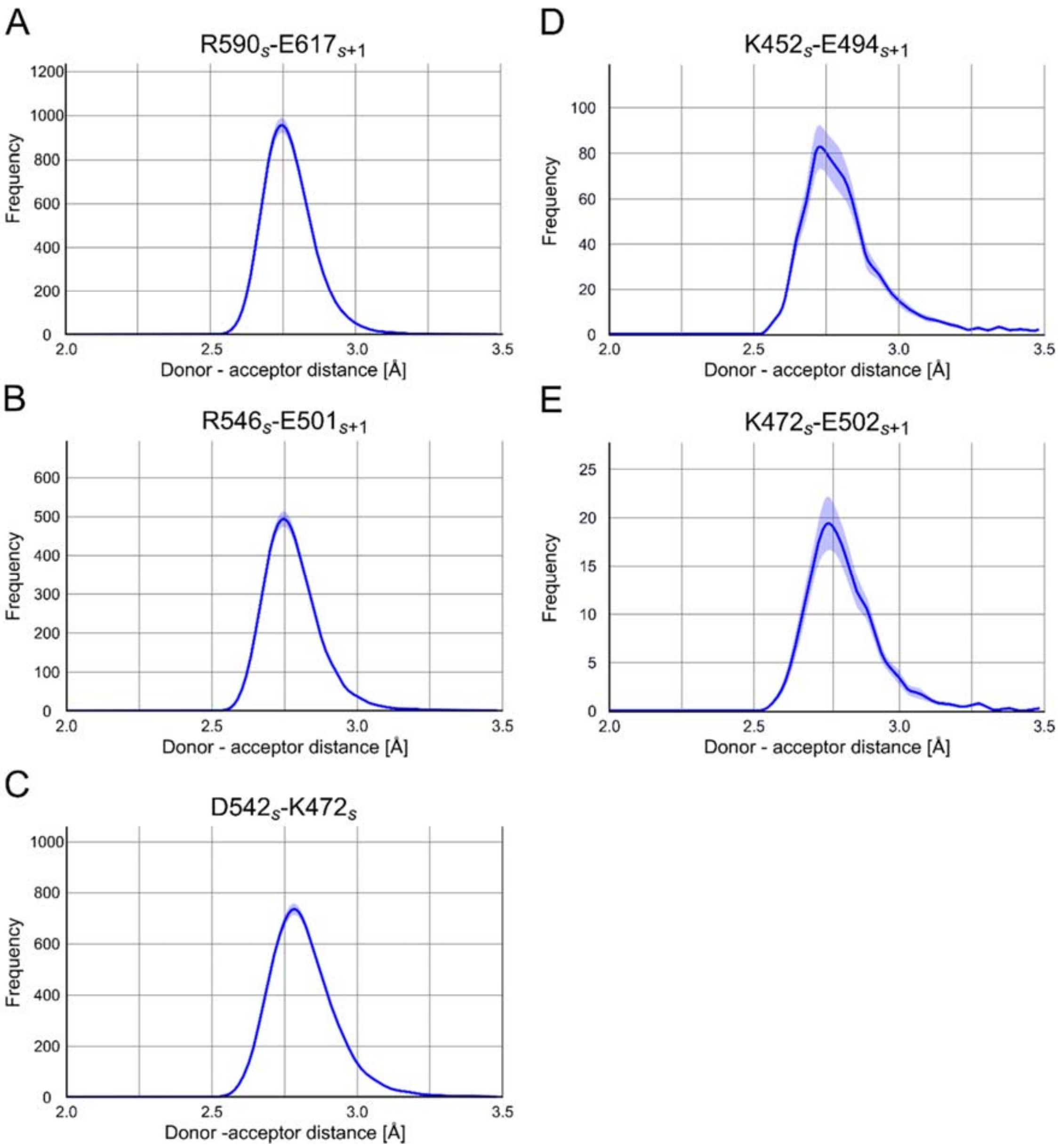


**Fig. S3.** Occurence frequency of inter- (A, B) and intra-subunit (C) salt bridges along predicted pathways from ten independent MD trajectories with in total 2,000 snapshots of the cAMP-bound CL-CNBD. In the case of more than one acceptor or donor atom on a sidechain, the respective shortest donor-acceptor distance was recorded. Shaded areas indicate the SEM. The index $s$ ($s$ + 1) denotes subunit $s$ ($s$ + 1). The two salt bridges K452$_{s}$-E494$_{s+1}$ (D) and K472$_{s}$-E502$_{s+1}$ (E) are reported in refs. (11, 12) to be functionally relevant for mHCN2 activation. However, both salt bridges only occur infrequently along the ten independent MD trajectories. They are not identified to be on a pathway by our perturbation approach.

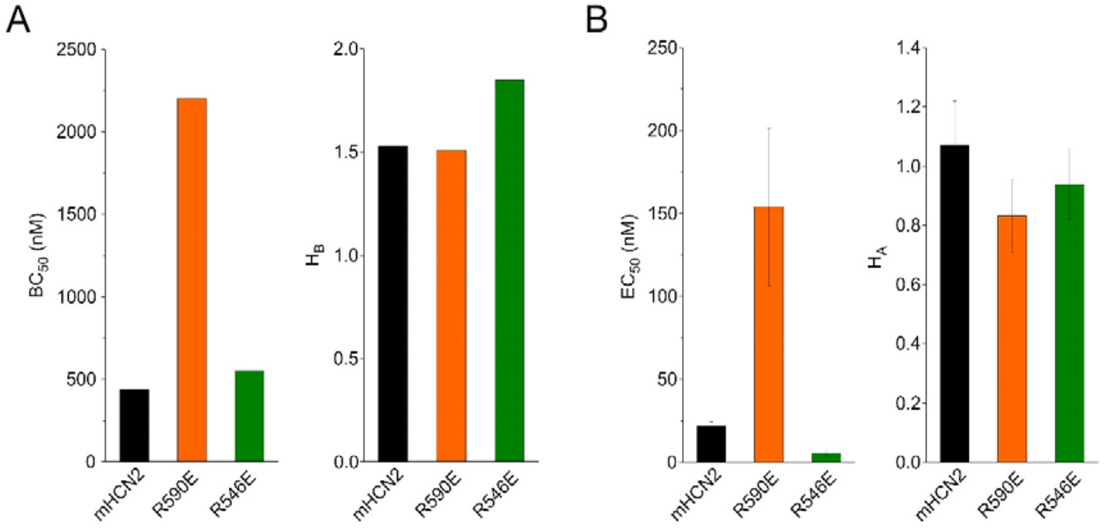


**Fig. S4.** Activation and binding after breaking the intersubunit salt bridges by reversing the charges in R590E and R546E. (A) Shown are the concentrations of half-maximum binding, $BC_{50}$ (left panel), and the Hill coefficients of binding, $H_B$ (right panel), for mHCN2, R590E, and R546E. Values were obtained from fitting the Hill equation (eq. 8 in the main text) to averaged relative fluorescence intensities (for details, see the Materials and Methods section and Fig. 1 in the main text). (B) Shown are the concentrations of half-maximum activation, $EC_{50}$ (left panel), and the Hill coefficients of activation, $H_A$ (right panel), for mHCN2, R590E, and R546E. Values were obtained from fitting the Hill equation (eq. 7 in the main text) to relative current amplitudes for individual recordings (for details, see the Materials and Methods section).

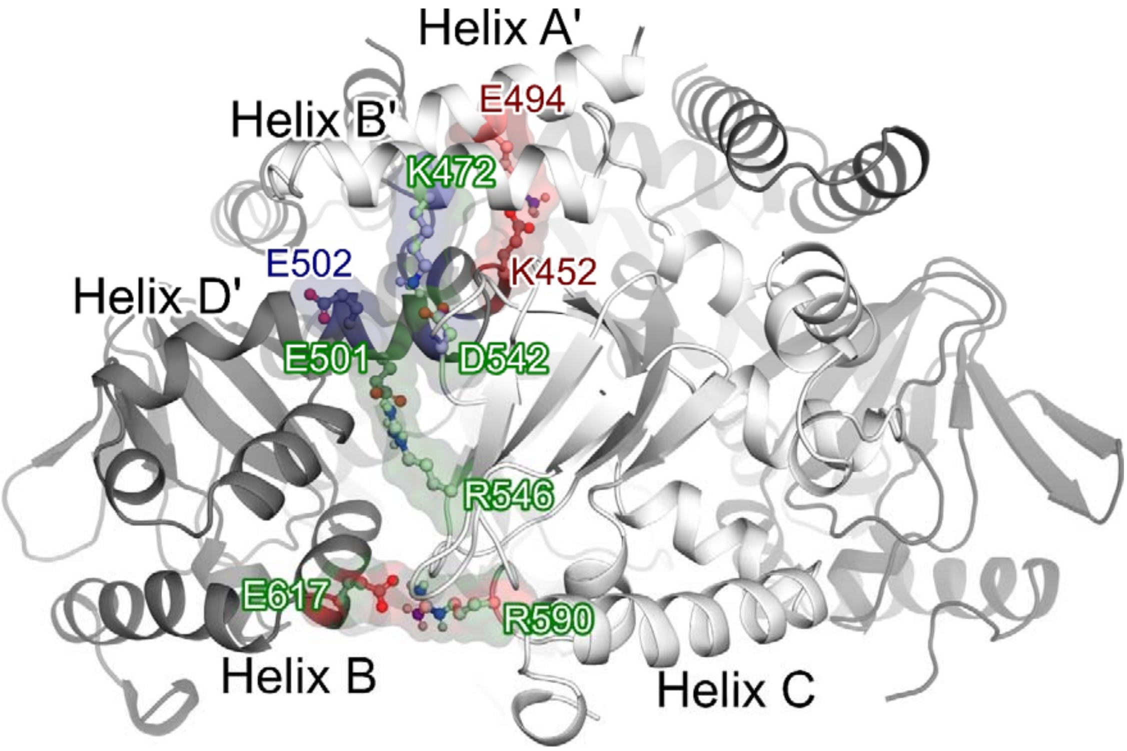


**Fig. S5:** Salt bridge interactions relevant in modulating cAMP affinity and potency. Surface representation in green depicts the $R546_{s}$-$E501_{s+1}$ interaction, herein described for the first time; representation in red shows the $K452_{s}$-$E494_{s+1}$ interaction reported by Marni et al. (11) and in blue the $K472_{s}$-$E502_{s+1}$ interaction reported by Craven and Zagotta (12); interactions shown in mixed colors indicate the $K472_{s}$-$D542_{s}$ (green-blue) and $R590_{s}$-$E617_{s+1}$ (green-red) salt bridges identified by us in agreement with results reported in ref. (12) and ref. (11).

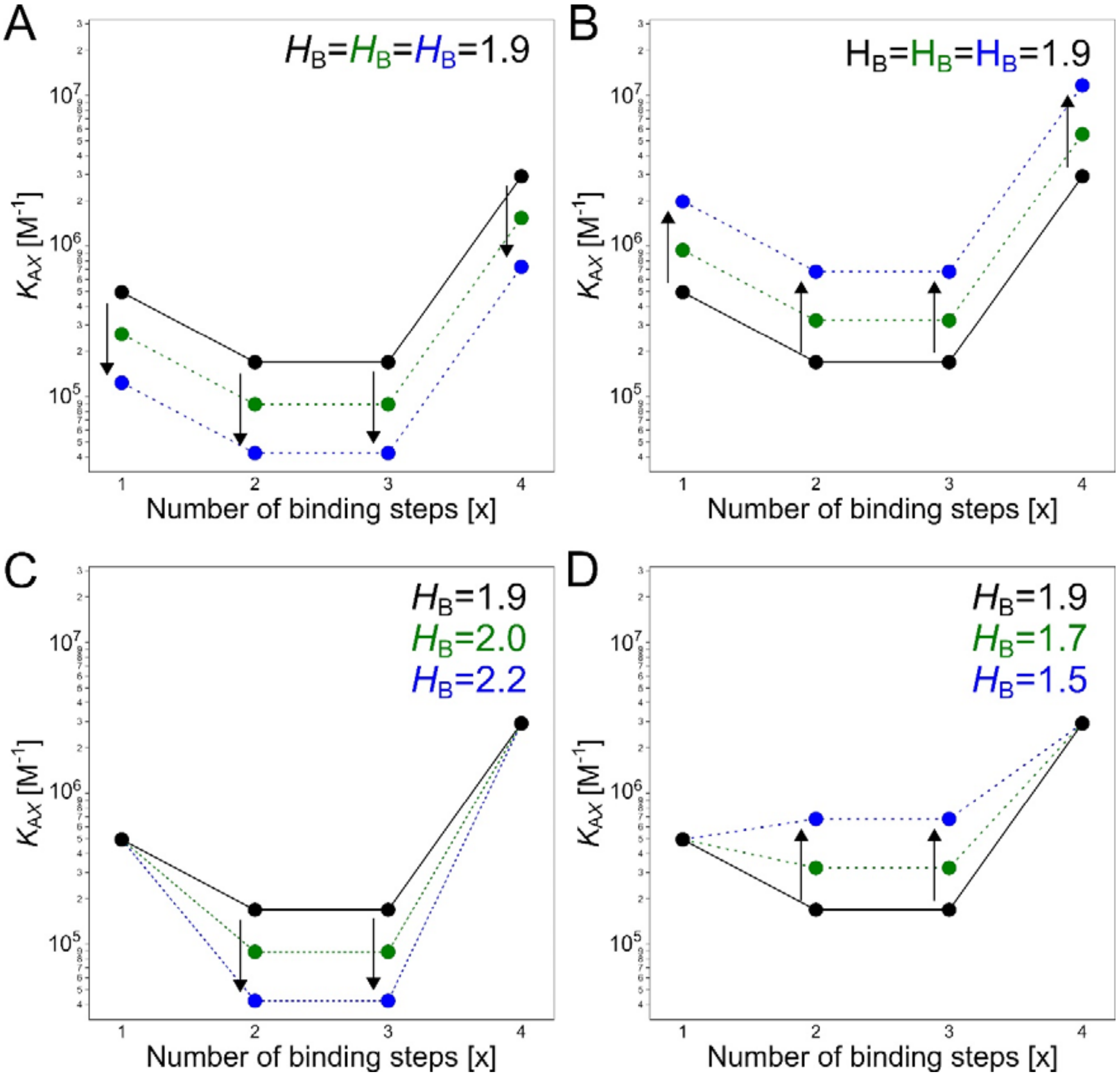


**Fig. S6.** Equilibrium association constants and how changing them affects the type of cooperativity and overall Hill coefficient $H_B$ (eq. S1), with $x$ is the number of binding steps. (A) Original equilibrium association constants are taken from ref. (2) and yield an overall Hill coefficient $H_B$ = 1.9 (eq. S1). Increasing (A) or lowering (B) all equilibrium association constants by a factor of 1.9 (green) and 4.0 (blue) does not change $H_B$. (C) Lowering the equilibrium association constants for the second and third binding step by a factor of 1.9 (4.0) results in $H_B$ = 2.0 (2.2). (D) Increasing the equilibrium association constants for the second and third binding step by a factor of 1.9 (4.0) results in $H_B$ = 1.7 (1.5).

**Supplementary Tables**

**Table S1. Sum of per-residue free energies associated with the change in structural stability due to the removal of (a) ligand(s) from the corresponding ligand-bound states.**

| Number of bund cAMP(s) | $\Delta\Delta G_{CNA}$[a, b] |
|---|---|
| *1* | 279.4 ± 0.7 |
| *2 (cis)* | 548.3 ± 1.8 |
| *2 (trans)* | 548.5 ± 4.7 |
| 3 | 839.0 ± 3.6 |
| 4 | 1185.0 ± 4.1 |

[a] Free energies in kcal $mol^{-1}$ computed from eq. 4 in the main text.
[b] Given are mean ± SEM ($n$ = 10).

**Table S2. Peristence of salt bridge interactions along 10 independent MD trajectories of the fully cAMP-bound state.**[a]

| Interaction[b] | Occurrence frequency [c] |
|---|---|
| $R590_s$-$E617_{s+1}$ | 70.5 ± 2.4 |
| $R546_s$-$E501_{s+1}$ | 39.1 ±1.6 |
| $D542_s$-$K472_s$ | 66.8 ± 4.0 |
| $K452_s$-$E494_{s+1}$ | 8.5 ± 0.9 |
| $K472_s$-$E502_{s+1}$ | 1.9±0.2 |

[a] A salt bridge interaction is present if the donor-acceptor distance is < 4 Å.
[b] The index $s$ ($s$ + 1) denotes subunit $s$ ($s$ + 1).
[c] Given are mean ± SEM ($n$ = 10) in %.

**Table S3. Activation and binding parameters for mHCN2 wildtype channels and mutated constructs.**[a]

| | HCN2 | R590A | R546A | K472A |
|---|---|---|---|---|
| $BC_{50}$ (nM) | 440 | 1780 | 640 | 232 |
| $H_B$ | 1.53 | 1.30 | 1.43 | 1.46 |
| $EC_{50}$ (nM) | 21.9 ± 2.59[b] | 206 ± 32.5[b] | 2.73 ± 0.50[b] | |
| $H_A$ | 1.07 ± 0.15[b] | 0.64 ± 0.04[b] | 1.15 ± 0.18[b] | |
| | **HCN2** | **R590E** | **R546E** | |
| $BC_{50}$ (nM) | 440 | 2200 | 550 | |
| $H_B$ | 1.53 | 1.51 | 1.85 | |
| $EC_{50}$ (nM) | 21.9 ± 2.59[b] | 154 ± 47.4[b] | 5.68 ± 1.65[b] | |
| $H_A$ | 1.07 ± 0.15[b] | 0.83 ± 0.12[b] | 0.94 ± 0.12[b] | |

[a] $BC_{50}$ is the concentration of half-maximum binding obtained from patch-clamp fluorometry recordings employing f1cAMP and non-activated channels (at -30 mV). $H_B$ is the Hill coefficient for binding. $EC_{50}$ is the concentration of half-maximum activation obtained from patch-clamp recordings employing untagged cAMP. $H_A$ is the Hill coefficient for activation.

[b] Given are mean ± SEM.

**Table S4 Important interactions for mHCN2 activation reported in the literature and predicted by us**

| Interaction[a] | Inter-subunit[b] | cNMP interaction[c] | Effect of breaking this interaction | Ref |
|---|---|---|---|---|
| K452$_{\text{Helix A'}}$-E494$_{\text{Helix C'}}$ | Yes | No | Abolishes cNMP effect on channel activation | (11) |
| K472$_{\text{Helix B'}}$-D542$_{\beta\text{-roll}}$ | No | No | Favors channel opening compared to wildtype; decreases the channel's apparent affinity for cAMP | (12) |
| K472$_{\text{Helix B'}}$-E502$_{\text{Helix D'}}$ | Yes | No | Favors channel opening compared to wildtype | (12) |
| R546$_{\beta\text{-roll}}$-E501$_{\text{Helix D'}}$ | Yes | No | Favors channel opening compared to the wildtype; modest effect on cAMP affinity | [d] |
| E582$_{\beta\text{-roll}}$- R632$_{\text{Helix C}}$ | No | No | Contributes to cAMP binding; modest effect on channel gating | (13) |
| R590$_{\beta\text{-roll}}$-E617$_{\text{Helix B}}$ | Yes | No | Impedes channel opening compared to wildtype; decreases the channel's apparent affinity for cAMP | (11) |
| E582$_{\beta\text{-roll}}$ | No | Yes | Contributes to cAMP binding; modest effect on channel gating | (13) |
| R591$_{\beta\text{-roll}}$ | No | Yes | Abolishes influence of cNMP on channel gating | (14) |
| T592$_{\beta\text{-roll}}$ | No | Yes | Decreases the channel's apparent affinity for cGMP while barely affecting cAMP affinity | (15) |
| R632$_{\text{Helix C}}$ | No | Yes | Abolishes cAMP influence | (13) |
| I636$_{\text{Helix C}}$ | No | Yes | Selectively stabilizes cAMP binding relative to cGMP binding; none of these residues contribute to channel gating | (13) |
| R635$_{\text{Helix C}}$ | No | No | Selectively stabilizes cAMP binding relative to cGMP binding; none of these residues contribute to channel gating | (13) |
| K638$_{\text{C-tail}}$ | No | No | Selectively stabilizes cAMP binding relative to cGMP binding; none of these residues contribute to channel gating | (13) |

[a] Underlined interactions indicate those salt bridges predicted by our computational analyses.
[b] Interacting residues are either in adjacent subunits ("Yes") or within one subunit ("No").
[c] Direct interaction between residues and cNMP.
[d] Herein investigated for the first time.
